# Impact of Bacterial Membrane Vesicles on Cellular Responses in *Leishmania amazonensis*-Infected Macrophages *In Vitro*

**DOI:** 10.1101/2025.07.08.663775

**Authors:** Pedro Henrique Gallo-Francisco, Genesy Pérez Jorge, Guilherme Augusto Sanches Roque, Marina Flóro e Silva, Natália Scanavachia da Silva, Cecília Costa Fagundes, Alexandre Leite Rodrigues Oliveira, Fausto Almeida, Marcelo Brocchi, Selma Giorgio

## Abstract

*Leishmania amazonensis* is an intracellular protozoan parasite and the main cause of Localized Cutaneous Leishmaniasis (LCL). A practical problem that can worsen the condition of infected individuals is secondary co-infection caused by opportunistic bacteria. This study evaluated the influence of Bacterial Membrane Vesicles (BMVs) from *Pseudomonas aeruginosa* and *Staphylococcus aureus*—both commonly associated with LCL—on macrophages previously infected with *L. amazonensis*. The diameter and concentration of BMVs were consistent with previous findings. We assessed infection dynamics, macrophage viability, and cytokine production. Results showed significant reductions in these parameters when *L. amazonensis-*infected macrophages were subsequently treated with BMVs, compared to the control group (infected but not treated with BMVs). Quantification of IL-2, IL-4, IL-6, IL-10, IL-17A, IFN-γ, and TNF-α revealed elevated levels of the last cytokine, suggesting an inflammatory response. Co-cultures treated with BMVs from *P. aeruginosa* (POMVs) exhibited a more pronounced inflammatory profile—marked by higher IL-1β and TNF-α concentrations—compared to those treated with BMVs from S. aureus (SEVs). IL-4 and IL-6 levels remained low relative to IL-1β and TNF-α. In conclusion, our data suggest that macrophage infection with *L. amazonensis* followed by exposure to bacterial MVs simulates a host-parasite-bacterial interaction, inducing strong immunogenic and inflammatory responses, which may represent potential targets for future vaccine strategies.

## Introduction

Leishmaniasis is an infectious disease caused by parasites of the genus *Leishmania* that represents a significant public health problem. There are about 20 species of parasites that are responsible for causing a broad spectrum of clinical conditions. Localized Cutaneous Leishmaniasis (LCL) is one of the most common forms of the disease, which manifests as single ulcerative lesions that can be self-resolving [1], [2], [3]. Typical LCL ulcers are painless and appear in exposed areas of the skin [4]. *L. amazonensis* causes LCL that responds to treatment or spontaneously heals, but can also lead to cutaneous dissemination. Leishmaniasis is endemic in 98 tropical and subtropical developing countries [3], [5].

The vectorial transmission of the parasites is made by dipterous psychodid sandflies belonging to the genus *Phlebotomus* in the Old World and by *Lutzomyia* in the New World [2], [4]. *Leishmania* parasites have a dixenic life cycle constituted by an extracellular flagellated promastigote and an intracellular amastigote form [2], [6]. The infectious metacyclic promastigotes are harbored in the female sandfly vector’s midgut and are transmitted to the vertebrate host during the blood meal [6]. Damage occurs to the dermis and its capillaries, leading to the recruitment of polymorphonuclear cells (PMNs), *e.g.,* neutrophils, dendritic cells, and macrophages are the first cells to migrate to infected sites, where the latter are the main host cell of *Leishmania* [2], [4]. After internalization, the phagosome and lysosome of macrophages fuse, forming a parasitophorous vacuole and the parasite transforms into the amastigote stage. Inside host cells, the parasite proliferates, breaks the membrane, and in the extracellular fluid infects naïve cells; alternatively, amastigotes surrounded by phagolysosomal components exit from damaged cells and infect other cells [2], [6]. In the next blood meal, the phlebotomine vector ingests host cells containing amastigotes that are transformed into metacyclic promastigotes in the midgut, where the parasite’s life cycle is complete [4], [6]. During the inoculation of the parasites, sandfly vectors insert their saw-like mouthparts into the skin [2], [6], [7], creating a phenomenon of indirect facilitation [8]. This event worsens the condition of vertebrate hosts, given that bacteria from both vertebrate host microbiota skin and invertebrate microbiota migrate into dermis, establishing a secondary co-infection inflammatory microenvironment [8].

In leishmaniasis, the polarization of the host’s immune response can influence the outcome of the disease, but genetic predisposition could be a factor. Mononuclear phagocytic cells, such as macrophages or neutrophils, can express different surface markers and a wide range of cytokines with discrete biological functions in the inflammatory microenvironment [9]. The activation pathways are described as classical activation (M1) and alternative activation (M2) [9]. The inflammatory M1-type activation profile may favor the host by eliminating intracellular LCL parasites, and this pattern of response leads to the differentiation of cells that secrete pro-inflammatory cytokines, such as IL-1β, IFN-γ, IL-12, and TNF-α [9]. While anti-inflammatory cytokines such as IL-4 or IL-10 promote M2-type activation profile [9].

During LCL secondary co-infections caused by opportunistic and/or pathogenic bacteria have been widely reported in the clinic [8], a fact that may worsen the condition of patients, associated with chronicity of lesions, prolonged hospitalization, and greater susceptibility to nosocomial infections [8], [10], [11]. The prevalence of secondary bacterial co-infections during LCL can vary between 18 - 80% and may be associated with the broad spectrum of skin lesions, clinical conditions, and immune response profiles [8]. Recent studies have even indicated that the bacteria-*Leishmania* co-infection occurs during the blood meal of the sandfly vector infected with *L. donovani*, where the vector exposes and deposits bacteria from its microbiota and the host, on the skin of mice [7].

Among the studies evaluated [8], the most identified bacteria from patients with LCL were the commensal *Staphylococcus aureus (S.a.)* and the opportunistic *Pseudomonas aeruginosa (P.a.),* bacteria that are commonly found in the healthy microbiota [8,12,13], which appear dysbiotic during LCL, that is, an imbalance in the microbial community compared to normal skin. Other studies have shown that *S.a.* is the microorganism most frequently identified from LCL, and it can exhibit different profiles of resistance to antibiotics used in the clinic [14], [15], [16].

In the context of secondary co-infection caused by bacteria during the LCL, the compromised epithelial tissue creates a gateway for opportunistic/pathogenic bacteria to colonize the inflammatory environment [8], which can secrete nanostructures described as Bacterial Membrane Vesicles (BMVs). BMVs are nanometric structures that are shaped spherically, varying between 20 and 400 nm in diameter [17], [18]. The BMVs are described as small granules on the surface of the bulged cell wall, which can be formed by evagination of the outer membrane in Gram-negatives, forming the outer membrane vesicles (OMVs) [19], [20]; or from the cytoplasmic content of Gram-positive bacteria, forming the extracellular vesicles (EVs), which do not have an outer membrane [21], [22]. Additionally, EVs can be secreted by all eukaryotic and prokaryotic cells, are the main mediators of extracellular communication, and carry protein, lipids, nucleic acids, and other active components that can be detected in body fluids [23].

The production of BMVs has been observed naturally [24], but also in tissues and fluids of infected mammalian cells, such as macrophages [25], [26], [27]. Macrophage-derived EVs play a critical role in mediating responses during infections and chronic inflammatory conditions [28], [29]. During *Leishmania* infection, macrophage-derived EVs present parasite antigens to dendritic cells, shaping adaptive immune responses and influencing disease outcomes [28]. Furthermore, *Leishmania*-derived EVs modulate host immune responses by facilitating cell-to-cell communication, delivering virulence factors, and enhancing parasite survival. Additionally, recent studies have highlighted the role of EVs in chronic inflammatory diseases, where they contribute to the persistence of inflammation by transferring pro-inflammatory mediators between cells [29], [30]. Also, EV-mediated communication between neutrophils and macrophages has been shown to promote anti-inflammatory metabolites, which help mitigate cytokine storm syndrome [30]. These findings underscore the dual role of EVs in both protective and pathological immune responses, making them key players in the regulation of immune processes and disease progression [28], [29], [30]. In the field of parasitology, research has explored EVs derived from other pathogens such as *Plasmodium* [31], [32], *Trypanosoma cruzi* [33], *Schistosoma* [34], *Trypanosoma brucei brucei* [35], and *Leishmania* [36]. As reported by Silverman and colleagues [28], EVs from *Leishmania* are capable of transporting virulence factors and altering macrophage activity, thereby promoting parasite survival through the suppression of pro-inflammatory cytokine secretion.

In this study, we aimed to evaluate the infection dynamics of RAW 264.7 macrophages infected with *L. amazonensis* and their subsequent interaction with BMVs derived from *S.a.* and *P. a.*, *in vitro.* These bacterial species are commonly found in skin lesions of patients with LCL and are associated with prolonged hospital stays, increased risk of nosocomial infections, and the need for additional therapeutic interventions. To mimic the presence of these clinically relevant bacteria, we used BMVs –specifically, OMVs from *P.a.* ATCC 27853 (POMVs) and EVs from *S.a.* ATCC 25923 (SEVs). We assessed whether exposure to these vesicles alters the infection profile and cytokine production of macrophages previously infected with *L. amazonensis*.

## Materials and Methods

### Bacterial strains and in vitro growing conditions

*P. aeruginosa* ATCC 27853 (*P.a.*), and *S. aureus* ATCC 25923 (*S.a.*) were grown aerobically at 37 °C on lysogeny broth (LB) agar and LB broth (Oxoid), prepared according to Sambrook and Russell, 2001 [37]. Bacterial isolates were cultured onto LB agar and incubated at 37 °C for 18h to obtain isolated colonies. Colonies grown on LB agar were then transferred into LB broth and grown for 18h at 37 °C at 150 rpm to make pre-inoculum (New Brunswick Scientific Innova 44R – Eppendorf). The following day, the bacterial culture was diluted 1:100 in fresh LB broth and allowed to grow until the exponential phase (OD_600_= 0,6 with approximately 1 x 10^8^ CFU/mL) under the same conditions. The bacterial strains utilized in this study were cryopreserved at -80

### Cell line

Murine macrophage cell line RAW 264.7 (ATCC# TIB-71, Rio de Janeiro Cell Bank – BCRJ, BRA) was cultured in RPMI-1640 media supplemented with 10% fetal bovine serum (FBS), penicillin (100 IU/mL) and streptomycin (100 µg/mL) (Vitrocell Embriolife), 2mM L-glutamine (Sigma-Aldrich), and 10mM HEPES (Calbiochem). The RAW 264.7 macrophages were cultured in 25cm² or 75cm² cell culture flasks in an incubator (Shel Lab) at 37 °C, 5% CO_2_, 21% O_2_, and balanced N_2_ [36].

### Parasite culture

Promastigotes of *L. amazonensis* strain M2269 (MHOM/BR/73/M2269) were maintained in vitro in RPMI-1640 medium supplemented with 10% FBS, 50µg/mL gentamicin, 2mM L-glutamine, and 10mM HEPES (Vitrocell Embriolife). The parasites were cultured in 25cm² or 75cm² cell culture flasks in a 26 °C incubator. For in vivo maintenance of *L. amazonensis* parasites, infections were performed in six-week-old female BALB/c mice obtained from the Multidisciplinary Center for Biological Investigation (CEMIB/UNICAMP), approved by the Ethics Committee for Animal Use (CEUA) under protocol 5904-1/2021, as described earlier in the laboratory [39].

### POMVs and SEVs isolation

The isolation of BMVs was performed as described by Batista and colleagues [40]. Briefly, pre-inoculum of both *P.a.* and *S.a.* cultivated during 18h in the same conditions described above, was diluted (1:100) in fresh LB broth at 37 °C at 150 rpm until the exponential phase for 20 h, which is known to be associated with high vesicle production [18], [22], [24]. The bacterial cultures were harvested by centrifugation at 5,000×g at 4 °C for 45 minutes. The supernatants were filtered through a 0.22 μm PES membrane filter (Merck, USA), and concentrated through ultrafiltration/diafiltration in a 100-kDa dialysis membrane (Merck, USA). The supernatants were centrifuged at 13,000×g at 4 °C for 15 min and filtered again through a 0.22 μm filter (Merck, USA). Later, the supernatants were ultracentrifuged (Beckman Coulter Optima L-90 K centrifuge) at 308,448×g at 4 °C for 1 h and 30 minutes. The purified sample of vesicles of both *P.a.* and *S.a.* was resuspended in a 10mM HEPES/0.85% NaCl buffer, filtered again in a 0.22 μm filter, and stored at -80 °C for further analysis.

### Nanoparticle Tracking Analysis (NTA) of the membrane vesicles

Size-distribution analysis and quantification of BMV preparations were performed on a NanoSight NS300 (Malvern Instruments). Purified BMVs were analyzed using a NanoSight NS300. Both scatter and capture settings such as focus, camera, and gain settings were optimized to make particle tracks visible. Then measurements were obtained in triplicate and analyzed using NanoSight software (Ver 3.2.16). The data on the sizes of BMVs are expressed as means ± SD of size distribution.

### Bradford assay of the POMVs and SEVs

The total protein content of POMVs and SEVs was determined using the Bradford assay, with bovine serum albumin (BSA) (Sigma-Aldrich) as a reference for the calibration curve [41]. Data on total protein concentration of POMVs and SEVs are expressed as means ± SD, and the assay was repeated at least three independent experiments.

### L. amazonensis-infected RAW 264.7 murine macrophages treated with POMVs or SEVs

2 x 10^5^ macrophages/well of RAW 264.7 macrophages were seeded in a 24-well plate with sterile round coverslips, in 1mL of complete RPMI-1640 medium. The cell co-cultures were incubated overnight at 37 °C, 5% CO_2_, 21% O_2_, and balanced N_2_ to allow macrophage adherence. Subsequently, the culture medium was removed and discarded, and the wells were washed once with PBS to remove cellular debris. The macrophage co-cultures were then incubated for 24 h with *L. amazonensis* at a ratio of 1:10 (number of macrophage: number of parasites, respectively). The plate was further incubated at 37 °C, 5% CO_2_, 21% O_2_, and balanced N_2_. Afterward, the supernatants were discarded, and the wells were washed once with PBS to remove remaining parasites. The POMVs were then inoculated for 24h at a macrophage-to-vesicle ratio of 1:1 (=0.03ng of protein/well), 1:10 (=0.26ng of protein/well), and 1:100 (=2.58ng of protein/well). The SEVs were inoculated for 24h at a macrophage-to-vesicle ratio of 1:1 (=1.50ng of protein/well), 1:10 (=0.150ng of protein/well), and 1:100 (=1.5ng of protein/well). Both POMVs and SEVs were inoculated in the wells containing previously infected macrophages with the parasite. We evaluated *L. amazonensis*-infected RAW 264.7 macrophages without BMVs as a control of co-cultures that were infected with the parasite and then treated with BMVs. Also, we evaluated uninfected macrophages treated with POMVs or SEVs for 24h in the same cell-to-vesicle ratio mentioned above, as a control of co-cultures infected with the parasite and treated with BMVs, and to assess whether the infection influences the viability. The wells received 1mL of complete RPMI-1640 medium. The plate was incubated for 24 hours at 37 °C, 5% CO_2_, 21% O_2_, and balanced N_2_. After 24 h of incubation, the supernatants from the cell co-cultures were collected and stored at −20 °C for subsequent quantification of cytokines. An aliquot of these supernatants was plated on LB agar to check for contamination in the co-culture supernatants and was incubated at 37 °C for 18 h. The coverslips in the plate wells were fixed with methanol for 10 minutes and stained with Giemsa to visualize the morphological appearance of macrophages. Adherent macrophages with a structurally preserved nucleus and cytoplasm were considered viable and were quantified in 40 representative fields to assess viability, the number of *L. amazonensis*-infected macrophages, and the number of intracellular amastigotes [42]. The percentage of infected macrophages was calculated as the ratio of the total macrophages counted, and the number of amastigotes per infected macrophage was calculated by dividing the total number of intracellular amastigotes by the total number of infected macrophages [42]. The infection index was calculated as the product of the percentage of infected macrophages and the number of amastigotes per infected macrophage [42]. All counts were made under a 1000x optical microscope. Three independent experiments were performed in triplicate. The data of the analyzed parameters are expressed as the means ± SD.

### Transmission electron microscopy (TEM) of the POMVs and SEVs

Thawed vesicles stored at -80 °C were prepared for electron microscopy analysis as described by Sarra and colleagues [43]. Samples for TEM examinations were prepared at room temperature by depositing 20μL of vesicle suspensions on a 300-mesh copper grid for electron microscopy covered by Formvar film. A negative staining was realized by the addition of 10 μL of 2% uranyl acetate (Sigma Aldrich) solution. The vesicles were examined under a Tecnai G2 Spirit BioTwin (FEI) operating at 80kV, in 68,000x magnification.

### Cytometric Bead Array (CBA)

Quantification of interleukin-2 (IL-2), IL-4, IL-6, IL-10, IL-17A, interferon-γ (IFN-γ), and tumor necrosis factor-α (TNF-α) was carried out to investigate the profile of immune response in this co-culture system, and whether the macrophages exhibit a profile of susceptibility or resistance to parasite infection. Freeze supernatants of the healthy or uninfected macrophages treated with BMVs, and cell co-cultures containing RAW 264.7 macrophages infected with *L. amazonensis* and treated with POMVs or SEVs, were used for cytokine quantification. The cytokines were quantified by flow cytometry (FACSCanto II system, BD Biosciences) using BD Cytometric Bead Array (CBA) mouse Th1/Th2/Th17 cytokine kit (CBA, BD Biosciences) according to the manufacturer’s instructions. FCAP Array, v. 3.0, software (Becton Dickinson) was used for data analysis. Results are expressed in pg/mL as means ± SD, based on standard concentration curves of each cytokine [41]. All samples were made in triplicate, and the experiment was repeated in at least three independent assays.

### Enzyme-Linked Immunosorbent Assay (ELISA)

Freeze supernatants of the healthy or uninfected macrophages treated with BMVs, and cell co-cultures containing RAW 264.7 macrophages infected with *L. amazonensis* and treated with POMVs or SEVs, were used to quantify IL-1β. Quantifications of IL-1β were performed using a Rat IL-1β/IL-1F2 DuoSet ELISA Kit (R&D Systems), and all procedures followed the manufacturer’s instructions. The IL-1β concentration is expressed in pg/mL as means ± SD, based on a standard curve as reference values [44]. All samples were made in triplicate, and the experiment was repeated in at least three independent assays.

### Statistical analysis

For the statistical analysis, media ± SD was calculated, and variance analysis (ANOVA) was performed between experimental situations, on the software GraphPad Prism® Ver. 9.0.2 (San Diego, CA, EUA). The statistical significance was verified through the t-Student test with Welch’s correction, when applicable, and considered acceptable when p≤ 0,05.

## Results

### Characterization of the POMVs and SEVs by NTA, TEM, and protein concentration

To mimic the presence of bacteria in *L. amazonensis*-infected RAW 264.7 macrophages, we isolated and analyzed the POMVs and SEVs secreted by *P.a.* and *S.a.,* respectively. Characterization of the vesicles by NTA, TEM, and Bradford assay are shown in Figure 1, and Figure 2.

**Figure 1.**
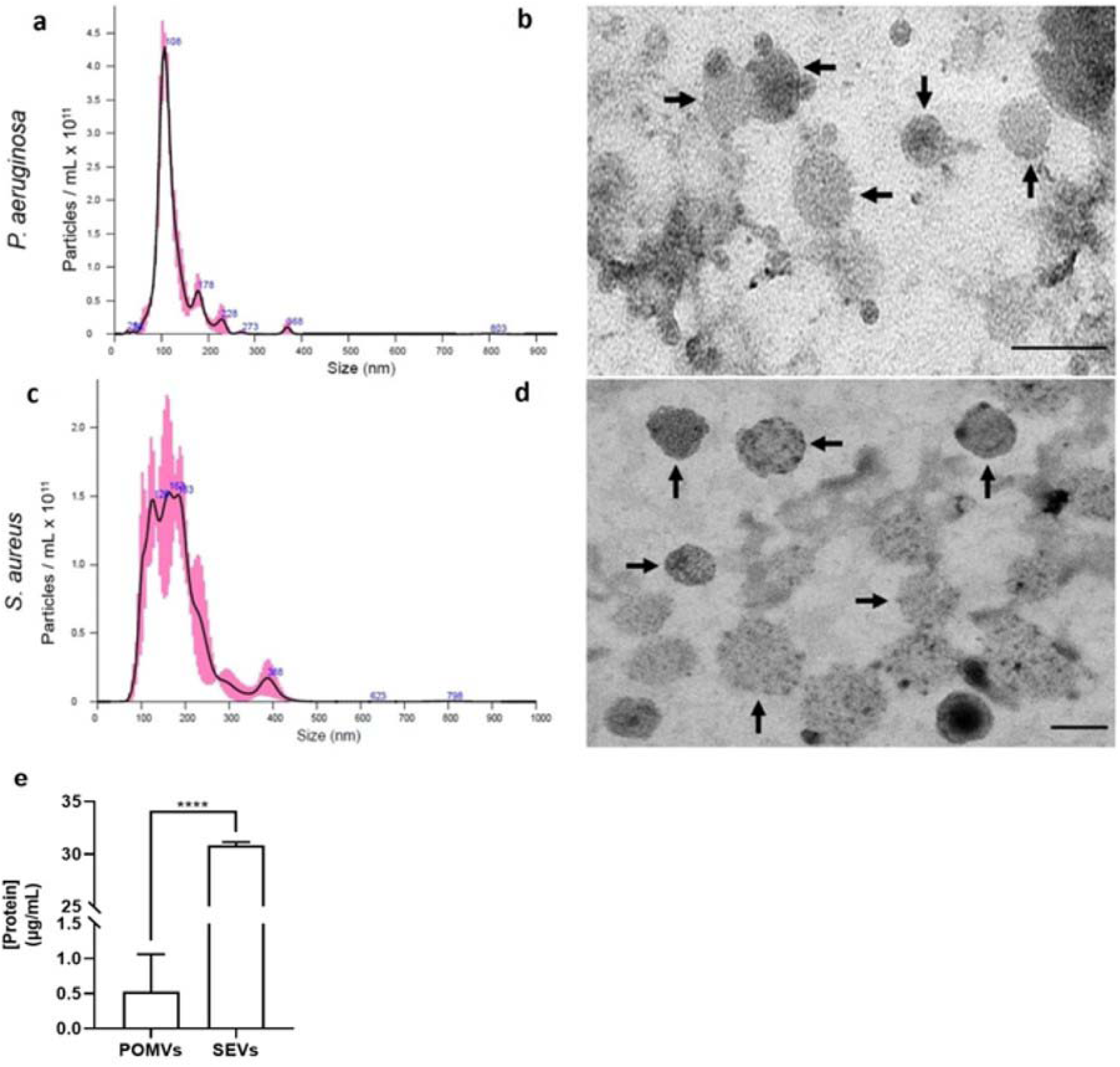
Characterization of the POMVs and SEVs by NTA, TEM, and Bradford assay. (a) and (c) NTA detects nanoparticles in solution making it possible to characterize vesicles in terms of diameter (in nm) and amount of particles/mL. (b) and (d) TEM of POMVs and SEVs. Black arrows indicate vesicles negatively stained with 2% uranyl acetate and visualized at 68.000x. Scale Bars = 100 nm. (e) Total protein quantification of POMVs and SEVs by Bradford assay. Data are shown as mean ± SD and are representative of three independent assays performed in triplicate. Statistical significance was calculated using the unpaired Student’s T-test. ****p≤ 0.0001.

**Figure 2.**
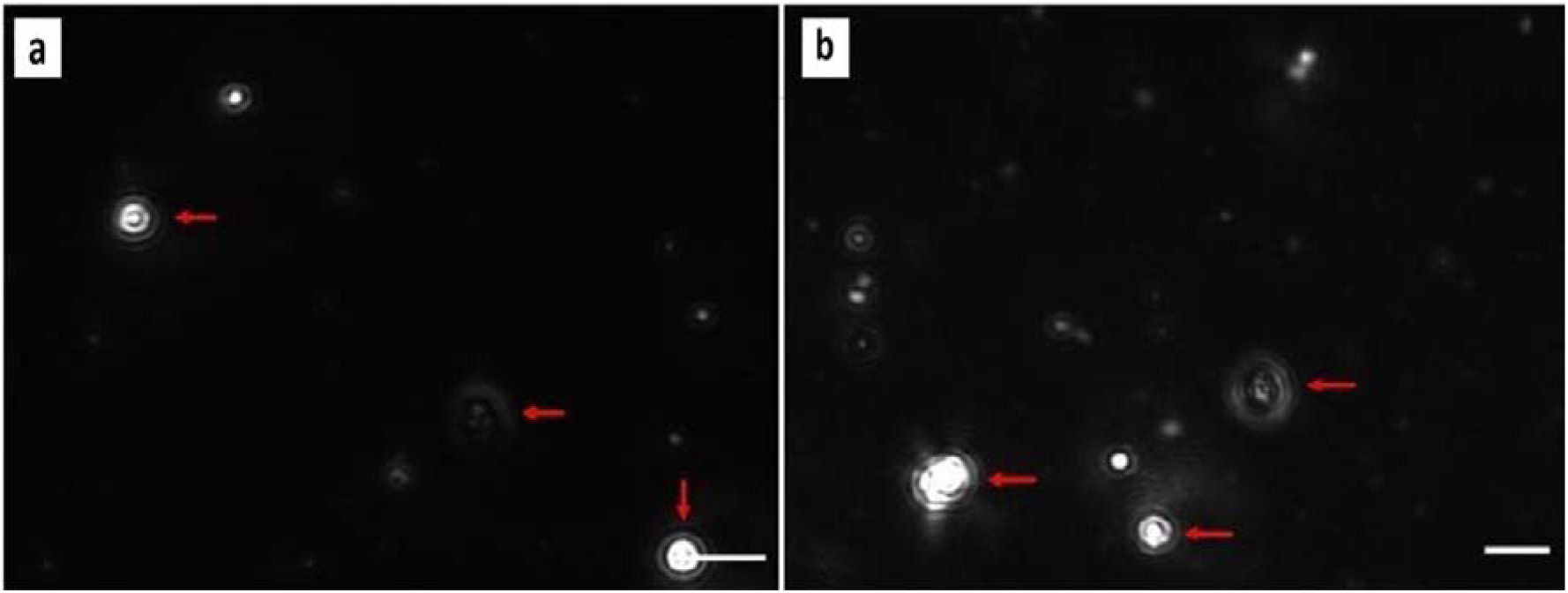
Capture of POMVs (a) and SEVs (b) observed under laser light scattering microscopy coupled to NTA. Red arrows indicate the presence of POMVs (a) and SEVs (b) in the diluted PBS suspension, visualized during determinations of diameter and number of nanoparticles. Scale Bars = 100 nm.

The NTA data of the POMVs revealed an average size of 124.1 ± 4.4 nm and an average concentration of 3.88 ± 0.39 x 10^11^ particles/mL (Figure 1a). Data reported in the literature showed similar sizes and concentrations of POMVs [18], [45], [46], [47]. SEVs revealed an average diameter of 180.6 ± 6.3 nm and an average amount of 4.15 ± 0.70 x 10^11^ particles/mL (Figure 1c). Akin to our results, literature findings demonstrated particle sizes and concentrations similar to those we obtained [21], [22], [48], [49].

POMVs and SEVs were observed by TEM to demonstrate their morphological structure, as well as confirm their diameter, as observed by the NTA, where it was possible to verify that the samples maintained their structural integrity during the extraction process. In Figures 1b and 1d, it is possible to observe POMVs and SEVs, respectively, negatively stained with 2% uranyl acetate and visualized at 68.000x magnification.

The total protein concentration of the POMVs was 0,53µg of protein/mL, significantly smaller when compared with the total protein concentration found in the SEVs, 30,83µg of protein/mL (Figure 1e). After NTA, TEM, and Bradford analyses, POMVs and SEVs were inoculated into macrophages.

### Viability and morphology of RAW 264.7 macrophages treated with POMVs or SEVs but not infected by L. amazonensis

Adherent macrophages with a structurally preserved nucleus and cytoplasm were considered viable cells in this study (Figure 3) [42]. Data showed that both POMVs and SEVs killed ∼25% and ∼36% respectively, of RAW 264.7 macrophages within 24h post-treatment with 1:100 ratio of cell to BMVs (Figure 3a), the highest amount tested. Furthermore, uninfected and untreated control macrophages, *i.e.*, those that were not treated with either POMVs or SEVs, remained with the cellular structure preserved and the nucleus and cytoplasm remained intact (Figure 3b) in comparison to the macrophages that were treated with POMVs and SEVs (Figure 3c - Figure 3h). Significant differences were observed in the viability of the macrophages between the uninfected and untreated control (considered as 100% of viability) and the rest of the experimental situations, containing macrophages treated for 24h with POMVs and SEVs (Figure 3a). Furthermore, significantly smaller differences were observed in the viability of RAW 264.7 macrophages between cell cultures treated with 1:1, 1:10, and 1:100 cell to SEV ratios, as seen in Figure 3a. The data suggest that SEVs killed more macrophages than POMVs (Figure 3a). For example, when RAW 264.7 macrophages were treated with 1:100 ratio of cell to SEVs, there was ∼36% killing, compared with ∼25% killing for macrophages treated with the same cell to POMV ratio (Figure 3a). Also, significant differences were found in the viability of the macrophages between cell cultures treated with 1:1 and 1:100 cell to SEVs ratios (Figure 3a). Macrophages untreated (Figure 3b) and treated with POMVs (Figures 3c – 3e) and SEVs (Figures 3f – 3h) were photographed in representative fields to evaluate their morphology. Cells without treatment with POMVs or SEVs exhibited macrophage-like morphology, indicated by the round or ameboid form and oval-shaped nucleus (Figure 3b). The macrophages treated with POMVs are more spread and larger (Figures 3c – 3e), and cells treated with SEVs underwent rounding and a small size but were still adherent (Figures 3f – 3h).

**Figure 3.**
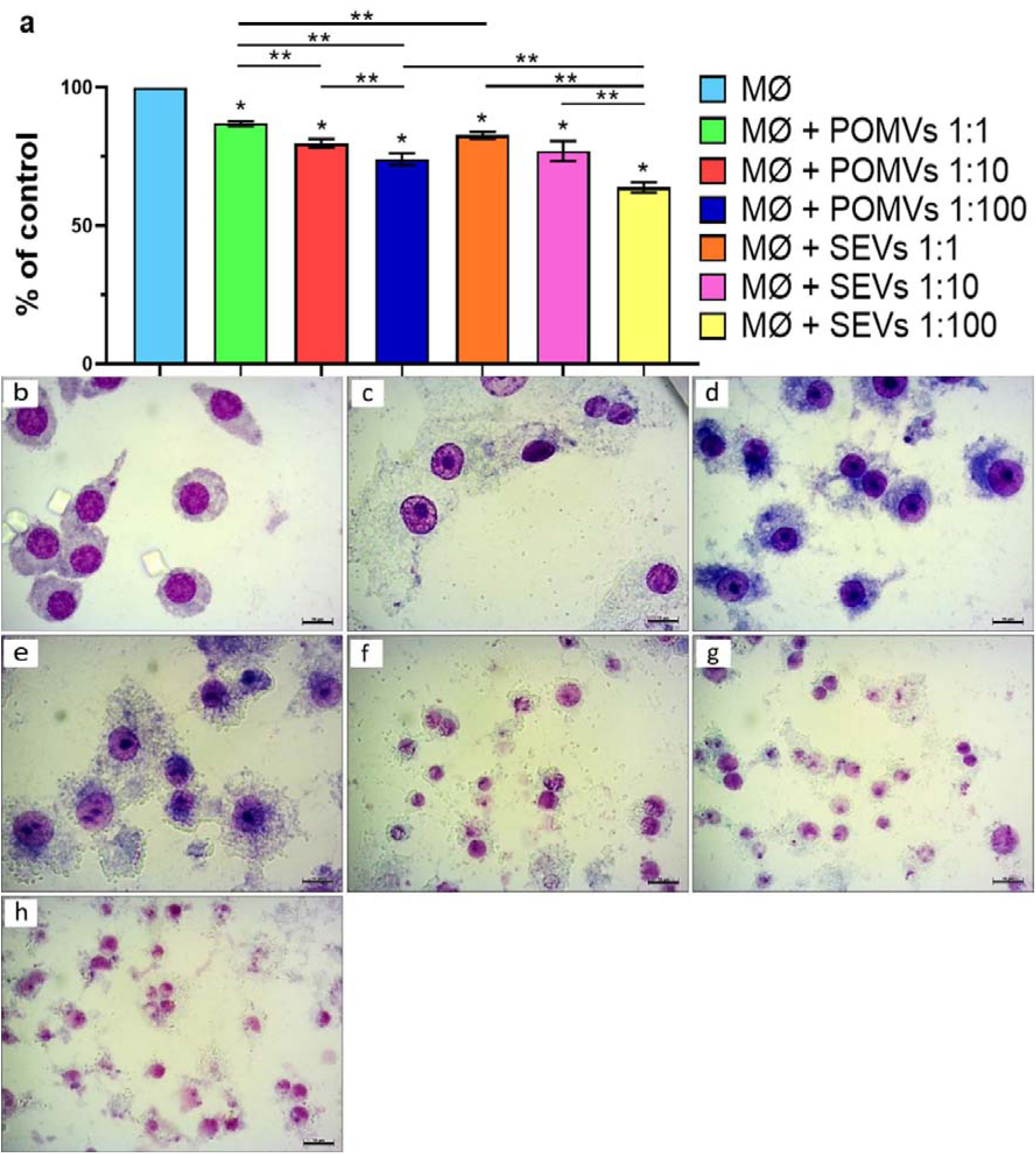
Adhesion and viability of macrophages from cell cultures without and with treatment with POMVs or SEVs; all without infection by the parasite. (a) macrophage viability, calculated as a percentage relative to the control. Data are shown as mean ± standard deviation and represent the results of three independent experiments performed in triplicate. MØ= uninfected macrophages with *L. amazonensis*, and untreated with POMVs or SEVs, as a control for the rest of the experimental situations; MØ + POMVs 1:1; MØ + POMVs 1:10; or MØ + POMVs 1:100= macrophages treated with 1:1 (=0.03ng of protein/well), 1:10 (=0.26ng of protein/well), or 1:100 (=2.58ng of protein/well) cell to POMVs ratios, respectively; MØ + SEVs 1:1; MØ + SEVs 1:10; or MØ’s + SEVs 1:100= macrophages treated with 1:1 (=1.50ng of protein/well), 1:10 (=0.150ng of protein/well), or 1:100 (=1.5ng of protein/well) cell to SEVs ratios, respectively. Macrophage cultures were treated with POMVs or SEVs for 24h and stained with Giemsa immediately afterward for microscopy quantification. Statistical significance was calculated using the unpaired Student’s T-test. *p≤ 0.05= significant difference between the control situation and the treatments. **p< 0.005= significant differences between treatments. (b) macrophages not treated with POMVs or SEVs; (c), (d), and (e) macrophages treated with 1:1, 1:10, and 1:100 cell to POMVs ratios, respectively; (f), (g), and (h) macrophages treated with 1:1, 1:10 and 1:100 cell to SEVs ratios, respectively. Macrophage cultures were treated with POMVs or SEVs for 24 h. Photomicrographs of representative fields, from three independent experiments carried out in triplicate, at 1000x magnification. Scale bars= 10μm.

L. amazonensis-infected RAW 264.7 macrophages and treated with POMVs or SEVs had a decrease in the infection index

The main objective of these assays was to investigate the influence of POMVs or SEVs treatment on RAW 264.7 macrophages previously infected with *L. amazonensis*. In the *in vitro* model we are using, infection of macrophages with *L. amazonensis* was established after 24h of incubation, followed by exposure to POMVs or SEVs for 24h, for analysis of the infection index and investigations into the modulation of the cellular immune response, i.e., production of cytokines. Macrophages infected with *L. amazonensis* come into contact with virulence factors isolated from bacteria, without direct contact between bacteria themselves, and consequently, destruction of the cell monolayer (Supplementary Figure 1). Initially, macrophages were quantified in experimental situations containing co-cultures of macrophages infected with *L. amazonensis* and untreated with POMVs or SEVs (control situation) and cell co-cultures infected with the parasite and treated with 1:1, 1:10, or 1:100 cell to POMVs or SEVs ratios (Figure 4).

**Figure 4.**
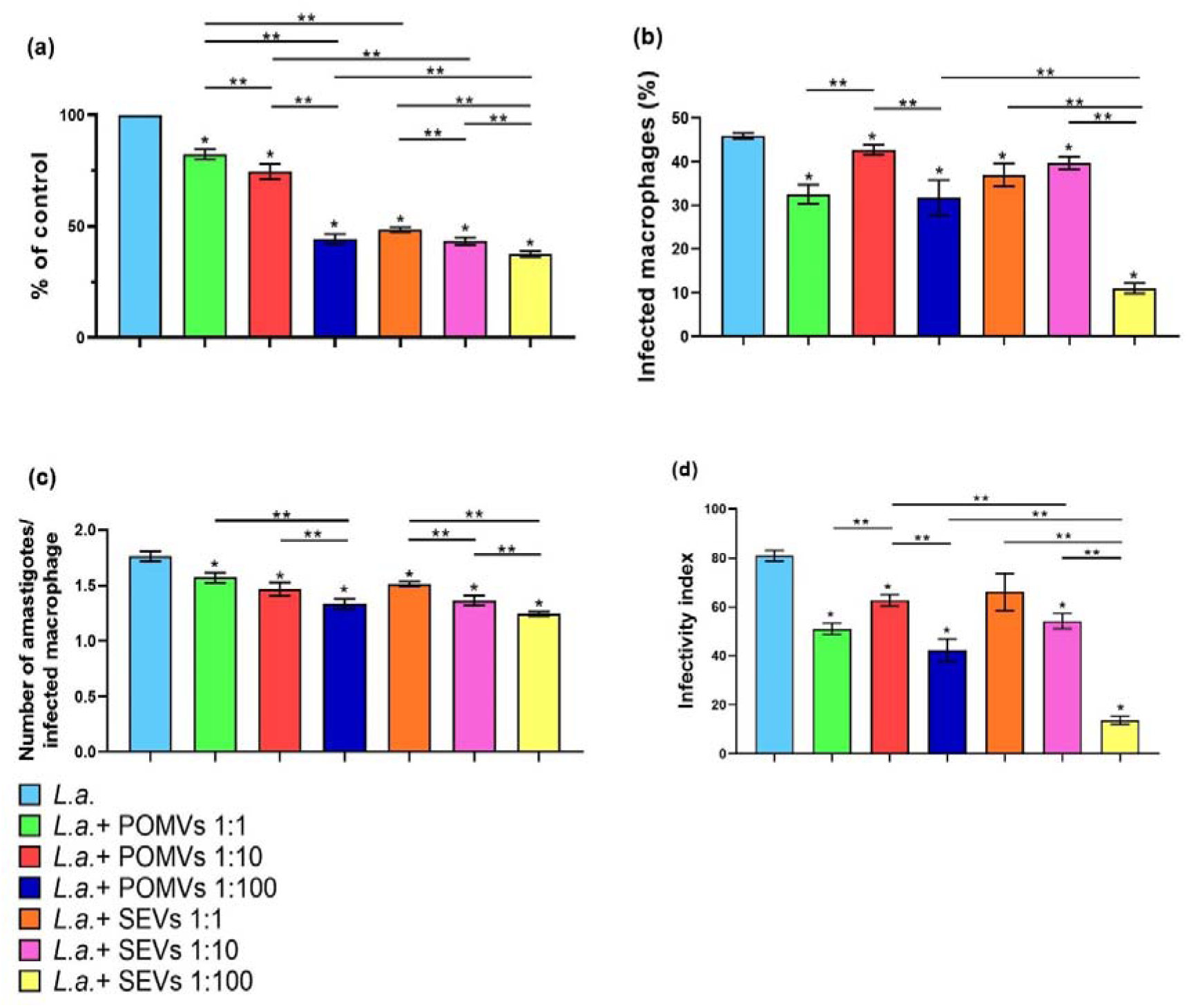
Adhesion, viability, and parasite load of RAW 264.7 macrophages, in cell co-cultures infected with *L. amazonensis* and treated with or not to POMVs or SEVs. (a) Macrophage viability, calculated as a percentage relative to the control, containing macrophages infected with *L. amazonensis*, and treated with POMVs or SEVs; (b) Average percentage of macrophages infected with *L. amazonensis* and treated with or not to POMVs or SEVs; (c) Number of amastigotes/ infected macrophage; (d) Infection rate of co-cultures of macrophages treated with or not to POMVs or SEVs. Data are shown as mean ± standard deviation and represent the results of three independent experiments performed in triplicate. *L.a.*= macrophages infected with *L. amazonensis* and untreated with POMVs or SEVs; *L.a.* + POMVs 1:1; *L.a.* + POMVs 1:10; or *L.a.* + POMVs 1:100= macrophages infected with *L. amazonensis* and treated with 1:1 (=0.03ng of protein/well), 1:10 (=0.26ng of protein/well), or 1:100 (=2.58ng of protein/well) cell to POMVs ratios, respectively; *L.a.* + SEVs 1:1; *L.a.* + SEVs 1:10; or *L.a.* + SEVs 1:100= macrophages infected with *L. amazonensis* and treated with 1:1 (=1.50ng of protein/well), 1:10(=0.150ng of protein/well) or 1:100 (=1.5ng of protein/well) cell to SEVs ratios, respectively. Macrophage co-cultures were infected with *L. amazonensis* for 24 h and then were treated with POMVs or SEVs for another 24 h. Statistical significance was calculated using the unpaired Student’s t-test. *p≤ 0.05= significant difference between the control situation, containing macrophages infected by the parasite without exposure to POMVs or SEVs, and the treatments; **p< 0.005= significant differences between treatments.

Significant differences were found in the viability of macrophages (Figure 4a), between the control situation and the rest of the treatments tested, suggesting a possible direct effect of the presence of POMVs or SEVs virulence factors on the viability of macrophages previously infected with the parasite. Significant differences were also observed in macrophage viability between cell co-cultures containing parasite infection and treatment with 1:1, 1:10 cell to POMVs ratios, and cell co-cultures infected by the parasite and treated with 1:100 cell to POMVs ratio; the data suggest a dose-dependent effect, between the decrease in macrophage viability and the increase in the proportion of POMVs tested (Figure 4a). Statistically smaller differences were found in macrophage viability between cell co-cultures treated with 1:1, 1:10, and 1:100 cell to SEVs ratios when compared with their corresponding counterparts containing cell co-cultures treated with 1:1, 1:10, and 1:100 cell to POMVs ratios (Figure 4a). Furthermore, the data suggested that SEVs compromised the viability of macrophages more than POMVs (Figure 4a). Data showed that both POMVs and SEVs killed ∼55% and 62% respectively, of RAW 264.7 macrophages within 24h post-treatment (Figure 4a) with 1:100 ratio of cell to BMVs. When macrophages were treated with 1:1, 1:10, and 1:100 cell to SEV ratios there was ∼51%, ∼56%, and ∼62% killing respectively, compared with ∼17%, ∼25%, and ∼55 killing of their corresponding counterparts containing cell co-cultures treated with 1:1, 1:10, and 1:100 cell to POMVs ratios (Figure 4a).

In Figure 4b, the average percentage of infected macrophages is observed, concerning the total number of infected macrophages counted. We noticed an evident significant decrease in the percentage of infected macrophages in all treatments tested. Significant decreases were observed between the average percentage of infected macrophages in the control situation, containing *L. amazonensis*-infected macrophages without treatment with POMVs or SEVs (45,92% of infected macrophages), and the remainder of all macrophage co-cultures infected with the parasite and subsequently treated with 1:1, 1:10, or 1:100 cell to POMVs or SEVs ratios (Figure 4b). A significant decrease was found between infected co-cultures treated with 1:100 cell to SEVs ratio and infected co-cultures treated with 1:100 cell to POMVs ratio (Figure 4b). The data suggest a positive effect of treatments with POMVs or SEVs on the intracellular parasites of the macrophages, probably caused by the vesicle’s virulence factors (Figure 4b).

After quantifying the intracellular amastigote forms, it was possible to observe significantly greater differences between the control situation and the rest of the treatments tested (Figure 4c). A significant decrease was found in the number of amastigotes per infected macrophage between cell co-cultures infected with the parasite and treated with 1:100 cell to POMVs ratio, and the following experimental situations: cell co-cultures infected with the parasite and treated with 1:1 and 1:10 cell to POMVs ratios (Figure 4c). A significant decrease was found between macrophage co-cultures infected with the parasite and treated with 1:100 cell to SEVs ratio, and the co-cultures infected with the parasite and treated with 1:1 cell to SEVs ratio (Figure 1c). These data suggest that significant reductions in the number of amastigotes per infected macrophage occurred due to the positive effect of treatment with 1:100 cell to POMVs or SEVs ratio on intracellular parasites (Figure 4c).

Regarding the infectivity index (Figure 4d), the product of the percentage of infected macrophages (Figure 4b), and the number of amastigotes/infected macrophage (Figure 4c), a significant reduction was found between the control co-cultures and the rest of the experimental situations (Figure 4d). These data suggest a trend towards a decrease in parasite load in parasite-infected co-cultures treated with POMVs or SEVs compared to the control without treatment with POMVs or SEVs (Figure 4d).

Significant decreases were found in the infection index between cell co-cultures infected with the parasite and treated with 1:10 and 1:100 cell to SEVs ratios when compared with their corresponding counterparts containing cell co-cultures infected with the parasite and treated with 1:10 and 1:100 cell to POMVs ratios (Figure 4d). These data suggest that the treatment with SEVs in the *L. amazonensis-*infected macrophages can decrease the parasite load due to virulence factors contained specifically and only in the SEVs, which are not present in the POMVs, and that, in some way, these virulence factors contained in SEVs may have had a positive effect in the intracellular parasites of macrophages treated with 1:10 and 1:100 cell to SEVs ratios (Figure 4d).

These experiments aimed to investigate the influence of POMVs or SEVs treatment on RAW 264.7 macrophages previously infected with *L. amazonensis*. Macrophages in representative fields were captured to highlight their morphological structure and the intracellular amastigote forms inside the cell host (data not shown). Interestingly, many cells with two nuclei were observed in these cell co-cultures (data not shown). Taken together, it can be concluded that the results (Figures 4a – 4d) indicated that the treatment with 1:100 cell to POMVs or SEVs ratios in the macrophage co-cultures compromises cell viability (Figure 4a), morphology of the cell monolayer (data not shown), which was directly altered by the virulence factors contained in the POMVs and SEVs, and infectivity index of macrophage co-cultures previously inoculated with *L. amazonensis* (Figure 4d).

### Cytokines induced by POMVs or SEVs in control cultures and L. amazonensis-infected RAW 264.7 macrophages

Cytokines IL-1β, IL-4, IL-6, and TNF-α were quantified after *L. amazonensis-*infected RAW 264.7 macrophages were treated with POMVs or SEVs, to investigate the profile of immune response in this co-culture system, and whether the macrophages exhibit a profile of susceptibility or resistance to parasite infection. Freeze supernatants from cell co-cultures were used for cytokine quantification.

Firstly, we analyzed the cytokine levels of the cell cultures treated with POMVs or SEVs, of the *L. amazonensis*-uninfected macrophages (Figures 5a – 5d).

**Figure 5.**
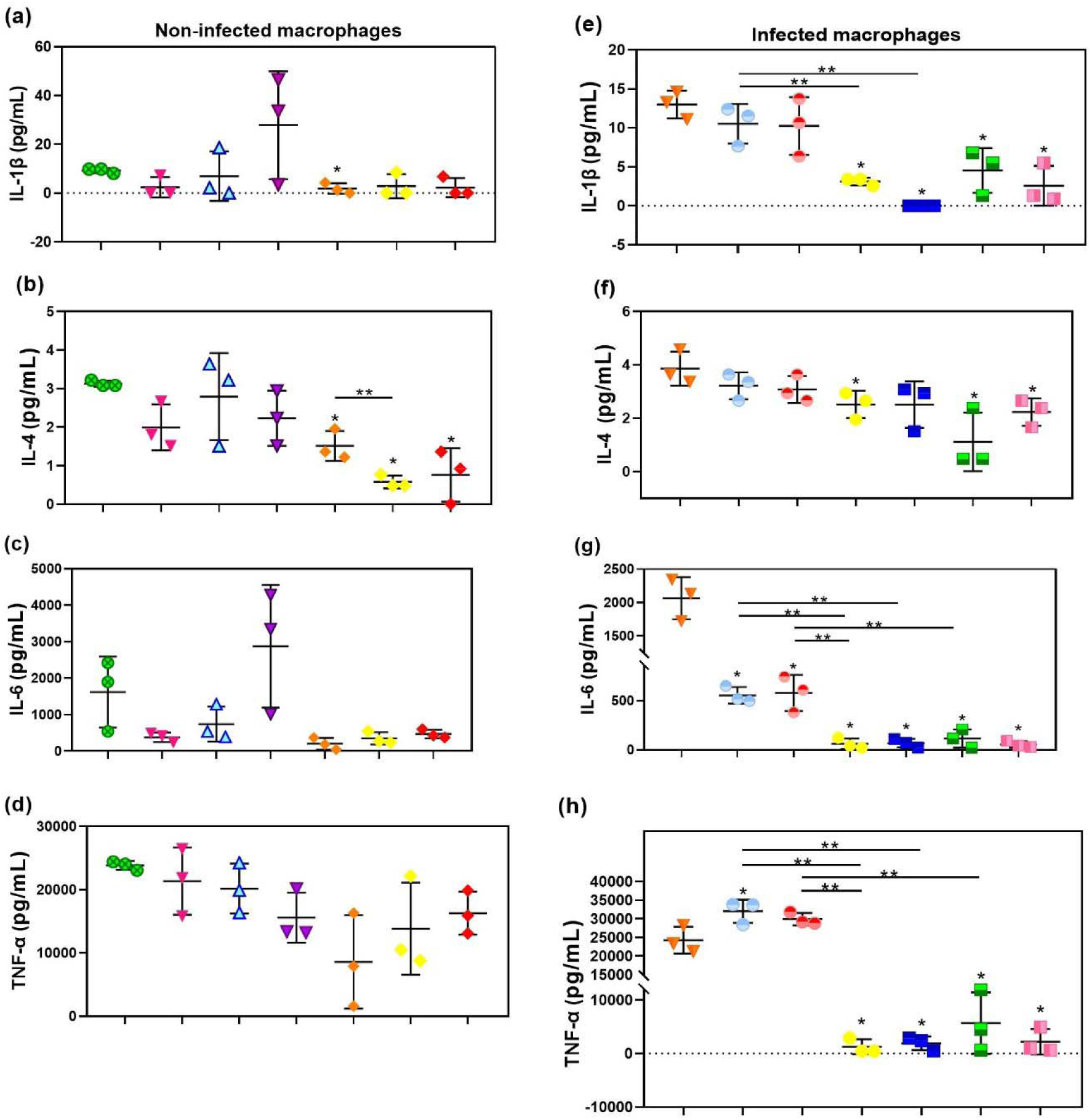
IL-1β, IL-4, IL-6, and TNF-α, in the supernatants of RAW 264.7 macrophages treated or untreated with POMVs or SEVs, and uninfected with the parasite (a – d); or in the supernatants of the cell co-cultures of *L. amazonensis*-infected RAW 264.7 macrophages treated or untreated with POMVs or SEVs (e – h). Data are shown as mean ± SD and represent the results of three independent assays performed in triplicate. (a – d): macrophages without infection by the parasite. 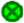= uninfected macrophages with *L. amazonensis*, and untreated with POMVs or SEVs, as a control for the rest of the experimental situations; 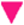= MØ + POMVs 1:1; 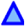= MØ + POMVs 1:10; 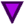= MØ + POMVs 1:100= macrophages uninfected with the parasite treated with 1:1 (=0.03ng of protein/well), 1:10 (=0.26ng of protein/well) and 1:100 (=2.58ng of protein/well) cell to POMVs ratios, respectively. 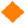= MØ’s + SEVs 1:1; 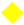= MØ’s + SEVs 1:10; 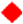= MØ’s + SEVs 1:100 = macrophages uninfected with the parasite treated with 1:1 (=1.50ng of protein/well), 1:10 (=0.150ng of protein/well) and 1:100 (=1.5ng of protein/well) cell to SEVs ratios, respectively. The macrophage cultures were treated with POMVs or SEVs for 24 h. (e – h): *L. amazonensis*-infected macrophages. 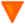 *L.a.*= macrophages infected with *L. amazonensis* untreated with POMVs or SEVs; 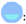= *L.a.* + POMVs 1:1; 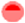= *L.a.* + POMVs 1:10; 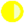= *L.a.* + POMVs 1:100= *L. amazonensis*-infected macrophages treated with 1:1 (=0.03ng of protein/well), 1:10 (=0.26ng of protein/well) and 1:100 (=2.58ng of protein/well) cell to POMVs ratios, respectively; 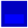 = *L.a.* + SEVs 1:1; 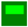 = *L.a.* + SEVs 1:10; 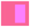 = *L.a.* + SEVs 1:100 = *L. amazonensis* infected macrophages treated with 1:1 (=1.50ng of protein/well), 1:10 (=0.150ng of protein/well) and 1:100 (=1.5ng of protein/well) cell to SEVs ratios, respectively. The macrophage co-cultures were infected with *L. amazonensis* for 24 h and after, were treated with POMVs or SEVs for 24 h. The statistical significance was calculated using the unpaired Student’s t-test. *p≤ 0,05= significative difference between the controls (uninfected macrophages with the parasite, and untreated with POMVs or SEVs (a – d); or *L. amazonensis*-infected macrophages without treatment with POMVs or SEVs (e – h)) and the rest of the experimental situations; **p< 0,005= significative difference between the treatments with POMVs or SEVs, for both uninfected macrophages (a – d), or *L. amazonensis*-infected macrophages.

The treatment of the macrophages with POMVs or SEVs resulted in altered concentrations of the assessed cytokines, *i.e.,* a significant reduction was found between the levels of IL-1β of the treated cultures when compared with the levels found in the untreated cell cultures with 1:1 cell to SEVs ratio (Figure 5a). Also, a significant decrease was found between the levels of IL-4 of the treated macrophages with POMVs or SEVs when compared with the concentrations found in the untreated cell cultures with 1:1, 1:10, and 1:100 cell to SEVs ratios (Figure 5b). A significant decrease was found in the levels of IL-4 in cell cultures treated with 1:10 cell to SEVs ratio when compared with the concentrations found in the cell cultures treated with 1:100 cell to SEVs ratio (Figure 5b). For both concentrations of IL-1β and IL-4, it was observed that secretion of these cytokines in the cell cultures treated with POMVs or SEVs was dramatically reduced when compared to the untreated control (Figures 5a and 5b). This may be due to the influence of BMVs on these cell cultures, which can suppress the production of these cytokines by macrophages, in some way (Figures 5a and 5b).

Regarding IL-6 and TNF-α levels in the cell cultures treated with 1:1, 1:10, or 1:100 cell to POMVs or SEVs ratios, no significant differences were found between the experimental situations (Figures 5c and 5d).

We also analyzed the cytokine levels of the *L. amazonensis-*infected RAW 264.7 macrophages treated with 1:1, 1:10, or 1:100 cell to POMVs or SEVs ratios for 24h (Figures 5e – 5h). A significant increase was found in the levels of IL-1β of the *L. amazonensis-*infected macrophages untreated with POMVs or SEVs and the following experimental situations: cell co-cultures infected with the parasite and treated with 1:100 cell to POMVs ratio; cell co-cultures infected with the parasite and treated with 1:1, 1:10 and 1:100 cell to SEVs ratios (Figure 5e). A significant increase was found in the levels of IL-1β of the cell co-cultures infected with the parasite and treated with 1:1 cell to POMVs ratio, and the following experimental situations: cell co-cultures infected with the parasite and treated with 1:100 cell to POMVs ratio (Figure 5e); and cell co-cultures infected with the parasite and treated with 1:1 cell to SEVs ratio (Figure 5e).

The IL-4 levels were significantly greater between *L. amazonensis-*infected RAW 264.7 macrophages untreated with POMVs or SEVs and the following experimental situations: cell co-cultures infected with the parasite and treated with 1:100 cell to POMVs ratio; and cell co-cultures infected with the parasite and treated with 1:10 and 1:100 cell to SEVs ratios (Figure 5f).

The IL-6 levels of macrophage co-cultures from the *L. amazonensis-*infected RAW 264.7 macrophages, were significantly greater when compared to the rest of the situations, containing cell co-cultures infected with the parasite and treated with 1:1, 1:10, and 1:100 cell to POMVs or SEVs ratios (Figure 5g). We observed significantly greater differences in the levels of IL-6 of the infected cell co-cultures treated with 1:1 and 1:10 cell to POMVs ratios when compared with their corresponding counterparts containing infected cell co-cultures treated with 1:1 and 1:10 cell to SEVs ratios (Figure 5g). Curiously, the IL-6 levels were significantly greater between infected cell co-cultures treated with 1:1 cell to POMVs ratio and infected cell co-cultures treated with 1:100 cell to POMVs ratio (Figure 5g). Also, statistically greater differences were found between infected cell co-cultures treated with 1:10 cell to POMVs ratio and infected cell co-cultures treated with 1:100 cell to POMVs ratio (Figure 5g).

Regarding TNF-α levels, significantly greater differences were found between *L. amazonensis-*infected RAW 264.7 macrophages, and the rest of the experimental situations, except for infected cell co-culture treated with 1:10 cell to POMVs ratio (Figure 5h). Significantly greater differences were found between infected cell co-cultures treated with 1:1 cell to POMVs ratio and infected cell co-cultures treated with 1:100 cell to POMVs ratio (Figure 5h). Statistically greater differences were found between infected cell co-cultures treated with 1:10 cell to POMVs ratio and infected cell co-cultures treated with 1:100 cell to POMVs ratio (Figure 5h). Significantly greater differences were found in the production of TNF-α in infected cell co-cultures treated with 1:1 and 1:10 cell to POMVs ratios when compared with their corresponding counterparts containing infected cell co-cultures and treatment with 1:1 and 1:10 cell to SEVs ratios (Figure 5h).

Overall, the data suggest that control macrophages and those infected with *L. amazonensis* exhibit distinct responses to POMVs or SEVs treatment, as evidenced by high production of IL-1β (Figures 5a and 5e) and TNF-α (Figures 5d and 5h) of both uninfected and infected macrophages. Nevertheless, our cytokine data suggested that infected cell co-cultures treated with POMVs exhibited significantly increased levels of IL-1β and TNF-α (Figures 5a and 5d, respectively), compared with infected cell co-cultures treated with SEVs. This profile of cytokine secretion observed could be due to the presence of virulence factors that the BMVs have, which trigger the production of inflammatory cytokines IL-1β (Figures 5a and 5e) and TNF-α (Figures 5d and 5h) by the RAW 264.7 macrophages. In contrast, IL-4 (Figures 5b and 5f) and IL-6 (Figures 5c and 5g) levels remained relatively low, suggesting a limited involvement of these cytokines in the response, compared with IL-1β and TNF-α.

## Discussion

Recent research has advanced our understanding of the microbiome in LCL lesions [8], [50], particularly the role of microorganisms and BMVs, which are immunogenic double-lipid membranes containing virulence factors [17], [51]. In this study, we isolated and purified POMVs and SEVs to explore whether the presence of these BMVs-containing virulence factors commonly observed in leishmaniotic lesions influences the persistence or elimination of the intracellular parasite *L. amazonensis,* during *in vitro* infection of the murine RAW 264.7 macrophages.

Our findings showed that POMVs averaged 124.1 ± 4.4 nm in size and an average of 3.88 ± 0.39 x 10¹¹ particles/mL (Figures 1a and 1b), consistent with previous data with similar strains that we used in this study, though slight variations in vesicle size and concentration were observed in another data of the literature, due to the use of different bacterial strains and culture conditions [18], [19], [20], [45], [46], [47], [52], [53], [54], [55]. Whereas SEVs averaged 180.6 ± 6.3 nm in diameter in our study and an average amount of 4.15 ± 0.70 x 10^11^ particles/mL (Figure 1c and 1d), aligning closely with prior findings [21], [22], [48], [49]. These comparisons underscore that the choice of bacterial strains and specific culture conditions significantly impact variations in the size, quantity of vesicles, and concentration of proteins [18].

After the characterization of POMVs and SEVs, experiments were performed with RAW 264.7 macrophages. These experiments were carried out to test the hypothesis proposed previously [8], where we suggested that the presence of bacteria during the LCL, as shown in literature reports, can alter the dynamics of *Leishmania* infection.

Our data showed that when RAW 264.7 macrophage cultures were treated with POMVs or SEVs for 24 h, in the absence of parasite infection, viability was not impaired, except for macrophage cultures treated with 1:1 cell to POMVs ratio, which showed significantly lower viability of the macrophages, compared to control cultures without treatment with vesicles (Figure 3a). Although viability was not affected, the morphological aspect of cells treated with POMVs and SEVs was altered (Figures 3c – 3h) compared to the morphology of control cells (Figure 3a). The macrophages treated with POMVs are more spread and larger, with an inflammatory appearance (Figures 3c – 3e) [56], and cells treated with SEVs underwent rounding and small size but were still adherents (Figures 3f – 3h). Interestingly, *L. amazonensis*-infected macrophages treated with POMVs or SEVs, had their viability amended and cellular morphology altered (Figure 4a); only macrophages infected with *L. amazonensis* and treated with POMVs had an inflammatory appearance (data not shown). Considering the data obtained with the quantification of total protein of POMVs and SEVs (Figure 1e), of which the highest concentration of total protein was found for SEVs, therefore, the cell cultures treated with SEVs containing more virulence factors with immunogenic and/or immunostimulatory properties, promoted a significant decrease in the viability of the macrophages, compared to the viability of the macrophages of the cell cultures treated with POMVs in the same proportions (Figure 4a). Taken together, the results indicated that the presence of BMVs in the macrophage cultures can alter both viability and morphology, although the infected cells appear to be more sensitive to this effect of the vesicles (Figure 3, Figure 4 and Figure 5). Several authors showed that OMVs have toxic effects on cells [50], and our findings (Figure 3, Figure 4 and Figure 5) showed that the mortality rate is similar to that of Deo and colleagues (2020). OMVs can activate pyroptosis [57], apoptosis [50], and inflammation [58]. For example, Deo and coworkers [50] showed that OMVs from non-pathogenic *E. coli* killed about 40% of murine bone marrow-derived macrophages as well as OMVs from *Neisseria gonorrhoeae* or *P.a.* The OMVs of *P.a.* diffuse into the host cell membrane and release virulence factors in a manner dependent on the actin filament pathway and, consequently, induce high cytotoxicity [59]. Data consistent with our findings were also observed with *Klebsiella pneumonia*-OMVs in activating the death mechanism in human bronchial epithelial cell lines [60]. Regarding previously *L. amazonensis*-infected RAW 264.7 macrophages treated with POMVs or SEVs, although there are no research articles dealing with the same cell co-culture system that we used in this work, it has been shown in other studies that the treatment of *Leishmania*-infected macrophages with IFN-γ or active parts of this biomolecule in association or not with LPS, can reduce the parasite load of macrophages [61], [62]. Our data shows the infectivity index of macrophages significantly decreased in cell co-cultures with *L. amazonensis* and vesicles when compared with the control, containing macrophages infected by the parasite and untreated with vesicles.

BMVs have been described as carrying LPS, glycerophospholipids, outer membrane proteins (OMP), lipid A, DNA, RNA, exotoxins, and proteases, which can be presented to competent cells of the immune system, trigger the activation of signals and stimulate microbicidal activities in macrophages [63]. Although the molecules contained in POMVs and the SEVs used in this work have not yet been identified, multiple molecules contained in these BMVs may exert activation on macrophages and reduce the parasite load.

The mechanism and biomolecules implicated in the dynamics of macrophage activation and intracellular killing of *Leishmania* are relatively well known and involve a few signaling pathways, such as NFkB (nuclear factor-kappaB) and the production of cytokines and microbicidal molecules [64]. Since in leishmaniasis, the inflammatory activation profile favors the host by eliminating intracellular parasites, the supernatants of previously *L.* amazonensis-infected RAW 264.7 macrophages, treated or not with POMVs or SEVs, were collected for cytokines quantification by CBA and ELISA (Figure 5). There is no data reported in the literature dealing with infection of cell co-cultures with the parasite and sequential exposure to POMVs or SEVs, as was done in our work.

Our cytokine production data suggested that infected cell co-cultures treated with POMVs exhibited increased levels of IL-1β and TNF-α (Figures 5a and 5d, respectively), compared with infected cell co-cultures and treatment with SEVs. Regarding the concentration of IL-4 and IL-6 (Figures 5b and 5c, respectively) secreted in the supernatants of the infected cell co-cultures, it was not found high concentrations compared with concentrations of IL-1β and TNF-α (Figures 5a and 5d, respectively). Studies have shown enhanced inflammatory cytokine production in BMV-treated cells. For example, when the J774A.1 macrophage strain was treated with EVs from *Rhodococcus equis*, IL-1β, IL-6, and TNF-α were significantly increased in the supernatants of cell cultures [65], and the murine bone marrow-derived macrophages treated with POMVs from the K strain of *P.a.* and dendritic cells treated with OMVs from *Salmonella enterica* [66] enhanced IL-1β concentration [65]. Notably, EVs from *L. shawi* and *L. guyanensis* are internalized by macrophages, where they modulate innate immune receptors, induce a mixed pro- and anti-inflammatory response, and promote MHC-I expression *in vitro* [67]. In addition, EVs from *L. donovani* contain parasite-derived molecules, that induced the transcription of M2 macrophage polarization markers (e.g., arginase 1, IL-10, and IL-4R). In contrast, EVs from *Salmonella Typhimurium* induced M1 polarization markers (e.g., iNOS, TNF-α, and IFN-γ), highlighting pathogen-specific modulation of EV composition and function [68].

In conclusion, while the *in vitro* assays employed in this study may not fully replicate the intricate microenvironment of leishmaniotic lesions, our experimental findings suggest a compelling hypothesis. The BMVs associated with secondary infections in LCL interact with the local population of *L. amazonensis*-infected macrophages, leading to compromised cellular homeostasis and a subsequent increase in inflammatory cytokine production (Figure 5). This altered microenvironment may contribute to the chronicity of the lesions, highlighting the critical role of bacterial interactions in disease progression [8], [12]. Taken together, our hypothesis [8] was that, *in vivo, Leishmania* parasites damage skin barriers during the blood meal and LCL of the phlebotomine vector, creating an entry point for bacteria and disrupting the skin microenvironment, causing dysbiosis (*i.e.,* changes in the skin microbiota). The result of this interaction is the subsequent competition between *Leishmania* and bacteria for host resources [8]. We transpose this hypothesis to our study and hypothesize from the results obtained with exposure of macrophages infected with *L. amazonensis* and subsequent treatment with POMVs or SEVs for 24h (Figure 4), and from the quantification of cytokines in the supernatants of the cell co-cultures (Figure 5). There may be neutral interaction, since neither the parasite nor the virulence factors contained in the BMVs of the *P.a.* and *S.a.* were eliminated by the RAW 264.7 macrophages. Still, we observed that there was an attempt by the macrophages to eliminate the intracellular parasites (Figure 4c), which although we observed significantly smaller decreases compared to the control without treatment with vesicles, it was a frustrating attempt, since there was no decrease in the level of total cleaning.

Looking ahead, we aim to investigate the detailed content of POMVs, SEVs, and *L. amazonensis* EVs, as well as macrophage-derived vesicles, through MALDI-ToF-MS and HPLC. These investigations will enhance our understanding of membrane vesicle composition and position them as crucial targets for developing innovative drug and vaccine platforms.

## Acknowledgments

The authors thank the unknown referees for the constructive review of this paper. Thanks to Prof. Dr. Eneida de Paula and the technicians for their support in the ultracentrifugation and NTA steps for the purification of BMVs, made at the Department of Biochemistry and Tissue Biology, Institute of Biology, UNICAMP. Thanks to Prof. Dr. Carlos Amilcar Parada and Dr. Catarine Massucato Nishijima for their support in plotting the IL-1β quantification data by ELISA, made at the Department of Structural and Functional Biology, Institute of Biology, UNICAMP. Thanks to the Central Laboratory of High-Performance Technologies in Life Sciences (LaCTAD) at UNICAMP and its team for providing flow cytometry assays.

## Author contributions

Pedro Henrique Gallo-Francisco: Investigation, Conceptualization, Methodology, Visualization, Writing – review & editing, Formal analysis. Genesy Pérez Jorge: Investigation, Conceptualization, Methodology, Visualization, Writing – review & editing, Formal analysis. Guilherme Augusto Sanches Roque: Investigation, Conceptualization, Methodology, Visualization, Writing – review & editing, Formal analysis. Marina Flóro e Silva: Investigation, Conceptualization, Methodology, Visualization, Writing – review & editing, Formal analysis. Natália Scanavachia da Silva: Investigation, Methodology, Resources. Cecília Costa Fagundes: Investigation, Methodology, Resources. Alexandre Leite Rodrigues de Oliveira: Investigation, Methodology, Resources. Fausto Almeida: Investigation, Methodology, Resources. Marcelo Brocchi: Investigation, Conceptualization, Methodology, Visualization, Writing – review & editing, Formal analysis. Selma Giorgio: Investigation, Conceptualization, Methodology, Visualization, Writing – review & editing, Formal analysis. All authors have read and approved of the final work. All authors have read and agreed to the published version of the manuscript.

---

**Supplementary Figure 1.**
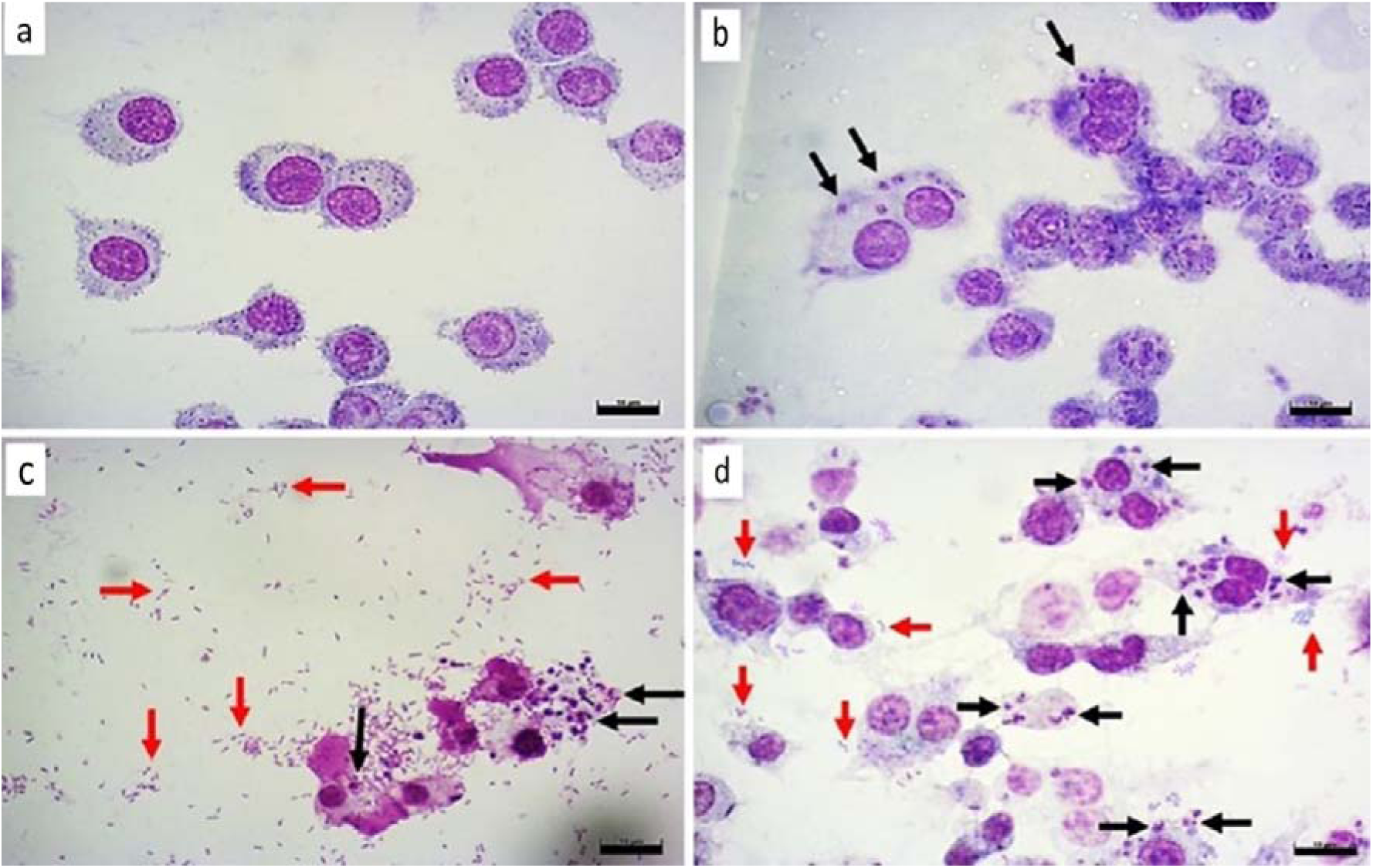
Photomicrographs of RAW 264.7 macrophages infected with *L. amazonensis* and inoculated or not to bacteria in the transwell system. Staining with Giemsa. (a) Control MØ = macrophages uninfected with *L. amazonensis*, and uninoculated with bacteria in the transwell inserts; (b) macrophages infected with *L. amazonensis* uninoculated with *P.a.* or *S.a.* in transwell inserts; (c) macrophages infected with *L. amazonensis* and inoculated with *P.a.* in transwell inserts; (d) macrophages infected with *L. amazonensis* and inoculated with *S.a.* bacteria in transwell inserts. Macrophage co-cultures were infected with *L. amazonensis* for 24 h, and after were inoculated with *P.a.* or *S.a.* in the transwell inserts for more 24 h. Black arrows indicate intracellular *L. amazonensis* in RAW 264.7 macrophages, and red arrows show *P.a.* or *S.a.* bacteria which translocated through the 0.4μm membrane of the transwell system. Photomicrographs captured in representative fields, from three independent experiments carried out in triplicate, at 1000x magnification. Scale bars= 10μm.

## References

[1] B. M. Scorza, E. M. Carvalho, and M. E. Wilson, “Cutaneous manifestations of human and murine leishmaniasis,” Jun. 18, 2017, MDPI AG. doi: 10.3390/ijms18061296.

[2] S. Burza, S. L. Croft, and M. Boelaert, “Leishmaniasis,” Sep. 15, 2018, Lancet Publishing Group. doi: 10.1016/S0140-6736(18)31204-2.

[3] World Health Organization (WHO)., “Leishmaniasis.”. May 20, 2024.

[4] B. Alemayehu and M. Alemayehu, “Leishmaniasis: A Review on Parasite, Vector and Reservoir Host,” Health Science Journal, vol. 11, no. 4, 2017, doi: 10.21767/1791-809x.1000519.

[5] E. Torres-Guerrero, M. R. Quintanilla-Cedillo, J. Ruiz-Esmenjaud, and R. Arenas, “Leishmaniasis: a review,” F1000Res, vol. 6, no. 750, pp. 1–15, May 2017, doi: 10.12688/f1000research.11120.1.

[6] Paul A Bates, “Revising Leishmania’s life cycle,” Nat Microbiol, vol. 3, no. 5, pp. 529–530, May 2018.

[7] R. Dey et al., “Gut Microbes Egested during Bites of Infected Sand Flies Augment Severity of Leishmaniasis via Inflammasome-Derived IL-1β,” Cell Host Microbe, vol. 23, no. 1, pp. 134–143.e6, Jan. 2018, doi: 10.1016/j.chom.2017.12.002.

[8] P. H. Gallo-Francisco, M. Brocchi, and S. Giorgio, “Leishmania and its relationships with bacteria,” Feb. 01, 2022, Future Medicine Ltd. doi: 10.2217/fmb-2021-0133.

[9] M. A. Hartley, S. Drexler, C. Ronet, S. M. Beverley, and N. Fasel, “The immunological, environmental, and phylogenetic perpetrators of metastatic leishmaniasis,” 2014, Elsevier Ltd. doi: 10.1016/j.pt.2014.05.006.

[10] D. Van Der Vliet et al., “Pseudomonas aeruginosa otochondritis complicating localized cutaneous leishmaniasis: Prevention of mutilation by early antibiotic therapy,” American Journal of Tropical Medicine and Hygiene, vol. 75, no. 2, pp. 270–272, 2006, doi: 10.4269/ajtmh.2006.75.270.

[11] D. Velozo et al., “Fatal mucosal leishmaniasis in a child * Leishmaniose mucosa fatal em criança *.”

[12] C. Gimblet et al., “Cutaneous Leishmaniasis Induces a Transmissible Dysbiotic Skin Microbiota that Promotes Skin Inflammation,” Cell Host Microbe, vol. 22, no. 1, pp. 13–24.e4, Jul. 2017, doi: 10.1016/j.chom.2017.06.006.

[13] A. Han et al., “The importance of a multifaceted approach to characterizing the microbial flora of chronic wounds,” Wound Repair and Regeneration, vol. 19, no. 5, pp. 532–541, Sep. 2011, doi: 10.1111/j.1524-475X.2011.00720.x.

[14] A. Patricia Isaac-Márquez and C. Manuel Lezama-Dávila, “Detection of Pathogenic Bacteria in Skin Lesions of Patients with Chiclero’s Ulcer. Reluctant Response to Antimonial Treatment,” 2003.

[15] L. A. Vera, J.L. Soares De Macedo, I. A. Ciuffo, G. Santos, and J. B. Santos, “Antimicrobial susceptibility of aerobic bacteria isolated from leishmaniotic ulcers in Corte de Pedra, BA,” 2006.

[16] L. de F. Antonio, et al., “Effect of secondary infection on epithelialisation and total healing of cutaneous leishmaniasis lesions,” Mem Inst Oswaldo Cruz, vol. 112, no. 9, pp. 640–646, Sep. 2017, doi: 10.1590/0074-02760160557.

[17] M. Toyofuku, N. Nomura, and L. Eberl, “Types and origins of bacterial membrane vesicles,” Jan. 01, 2019, Nature Publishing Group. doi: 10.1038/s41579-018-0112-2.

[18] N. J. Bitto, L. Zavan, E. L. Johnston, T. P. Stinear, A. F. Hill, and M. Kaparakis-Liaskos, “Considerations for the Analysis of Bacterial Membrane Vesicles: Methods of Vesicle Production and Quantification Can Influence Biological and Experimental Outcomes,” 2021. [Online]. Available: https://journals.asm.org/journal/spectrum

[19] D. Augustyniak, T. Olszak, and Z. Drulis-Kawa, “Outer Membrane Vesicles (OMVs) of Pseudomonas aeruginosa Provide Passive Resistance but Not Sensitization to LPS-Specific Phages,” Viruses, vol. 14, no. 1, Jan. 2022, doi: 10.3390/v14010121.

[20] V. L. D. Siqueira et al., “Structural changes and differentially expressed genes in Pseudomonas aeruginosa exposed to meropenem-ciprofloxacin combination,” Antimicrob Agents Chemother, vol. 58, no. 7, pp. 3957–3967, 2014, doi: 10.1128/AAC.02584-13.

[21] E. Y. Lee et al., “Gram-positive bacteria produce membrane vesicles: Proteomics-based characterization of Staphylococcus aureus-derived membrane vesicles,” Proteomics, vol. 9, no. 24, pp. 5425–5436, Dec. 2009, doi: 10.1002/pmic.200900338.

[22] M. Gurung et al., “Staphylococcus aureus produces membrane-derived vesicles that induce host cell death,” PLoS One, vol. 6, no. 11, Nov. 2011, doi: 10.1371/journal.pone.0027958.

[23] C. He, S. Zheng, Y. Luo, and B. Wang, “Exosome Theranostics: Biology and Translational Medicine,” Theranostics, vol. 8, no. 1, pp. 237–255, 2018, doi: 10.7150/thno.21945.

[24] N. Soler, M. Krupovic, E. Marguet, and P. Forterre, “Membrane vesicles in natural environments: a major challenge in viral ecology,” ISME J, vol. 9, no. 4, pp. 793–796, Mar. 2015, doi: 10.1038/ISMEJ.2014.184.

[25] J. D. Cecil et al., “Outer Membrane Vesicles Prime and Activate Macrophage Inflammasomes and Cytokine Secretion In Vitro and In Vivo,” Front Immunol, vol. 8, Aug. 2017, doi: 10.3389/fimmu.2017.01017.

[26] R. Finethy et al., “Inflammasome Activation by Bacterial Outer Membrane Vesicles Requires Guanylate Binding Proteins,” mBio, vol. 8, no. 5, Nov. 2017, doi: 10.1128/mBio.01188-17.

[27] M. L. Elizagaray et al., “Canonical and Non-canonical Inflammasome Activation by Outer Membrane Vesicles Derived From Bordetella pertussis,” Front Immunol, vol. 11, Aug. 2020, doi: 10.3389/fimmu.2020.01879.

[28] J. M. Silverman et al., “Leishmania Exosomes Modulate Innate and Adaptive Immune Responses through Effects on Monocytes and Dendritic Cells,” The Journal of Immunology, vol. 185, no. 9, pp. 5011–5022, Nov. 2010, doi: 10.4049/jimmunol.1000541.

[29] W. Ortmann, A. Such, I. Cichon, M. Baj-Krzyworzeka, K. Weglarczyk, and E. Kolaczkowska, “Large extracellular vesicle (EV) and neutrophil extracellular trap (NET) interaction captured in vivo during systemic inflammation,” Sci Rep, vol. 14, no. 1, Dec. 2024, doi: 10.1038/S41598-024-55081-X.

[30] H. Kang, T. Liu, Y. Wang, W. Bai, Y. Luo, and J. Wang, “Neutrophil–macrophage communication via extracellular vesicle transfer promotes itaconate accumulation and ameliorates cytokine storm syndrome,” Cell Mol Immunol, vol. 21, no. 7, pp. 689–706, May 2024, doi: 10.1038/s41423-024-01174-6.

[31] N. G. Sampaio, L. Cheng, and E. M. Eriksson, “The role of extracellular vesicles in malaria biology and pathogenesis,” Malar J, vol. 16, no. 1, p. 245, Dec. 2017, doi: 10.1186/s12936-017-1891-z.

[32] M. Gualdrón-López et al., “Characterization of Plasmodium vivax Proteins in Plasma-Derived Exosomes From Malaria-Infected Liver-Chimeric Humanized Mice,” Front Microbiol, vol. 9, Jun. 2018, doi: 10.3389/fmicb.2018.01271.

[33] L. M. de Pablos Torró, L. Retana Moreira, and A. Osuna, “Extracellular Vesicles in Chagas Disease: A New Passenger for an Old Disease,” Front Microbiol, vol. 9, Jun. 2018, doi: 10.3389/fmicb.2018.01190.

[34] G. G. Mekonnen et al., “Schistosoma haematobium Extracellular Vesicle Proteins Confer Protection in a Heterologous Model of Schistosomiasis,” Vaccines (Basel), vol. 8, no. 3, p. 416, Jul. 2020, doi: 10.3390/vaccines8030416.

[35] T. Dias-Guerreiro et al., “African Trypanosomiasis: Extracellular Vesicles Shed by Trypanosoma brucei brucei Manipulate Host Mononuclear Cells,” Biomedicines, vol. 9, no. 8, p. 1056, Aug. 2021, doi: 10.3390/biomedicines9081056.

[36] L. E. Soto-Serna et al., “Leishmania mexicana: Novel Insights of Immune Modulation through Amastigote Exosomes,” J Immunol Res, vol. 2020, pp. 1–12, Dec. 2020, doi: 10.1155/2020/8894549.

[37] J. Sambrook and D. Russell, Molecular Cloning: A Laboratory Manual, 3rd ed., vol. Vols 1, 2, 3. New York, 2001.

[38] A. Degrossoli, M. C. Bosetto, C. B. C. Lima, and S. Giorgio, “Expression of hypoxia-inducible factor 1alpha in mononuclear phagocytes infected with Leishmania amazonensis,” Immunol Lett, vol. 114, no. 2, pp. 119–125, Dec. 2007, doi: 10.1016/J.IMLET.2007.09.009.

[39] A. P. Araujo and S. Giorgio, “Immunohistochemical evidence of stress and inflammatory markers in mouse models of cutaneous leishmaniosis,” Arch Dermatol Res, vol. 307, no. 8, pp. 671–682, Oct. 2015, doi: 10.1007/S00403-015-1564-0/FIGURES/5.

[40] J. H. Batista et al., “Interplay between two quorum sensing-regulated pathways, violacein biosynthesis and VacJ/Yrb, dictates outer membrane vesicle biogenesis in Chromobacterium violaceum,” Environ Microbiol, vol. 22, no. 6, pp. 2432–2442, Jun. 2020, doi: 10.1111/1462-2920.15033.

[41] N. E. Zorgi et al., “Leishmania infantum transfected with toxic plasmid induces protection in mice infected with wild type L. infantum or L. amazonensis,” Mol Immunol, vol. 127, pp. 95– 106, Nov. 2020, doi: 10.1016/J.MOLIMM.2020.08.006.

[42] A. de Mesquita Barbosa, S. dos Santos Costa, J. R. da Rocha, C. A. Montanari, and S. Giorgio, “Evaluation of the leishmanicidal and cytotoxic effects of inhibitors for microorganism metabolic pathway enzymes,” Biomedicine & Pharmacotherapy, vol. 74, pp. 95–100, Aug. 2015, doi: 10.1016/j.biopha.2015.07.040.

[43] A. Sarra et al., “Biophysical Characterization of Membrane Phase Transition Profiles for the Discrimination of Outer Membrane Vesicles (OMVs) From Escherichia coli Grown at Different Temperatures,” Front Microbiol, vol. 11, Feb. 2020, doi: 10.3389/fmicb.2020.00290.

[44] R. A. Herman, P. N. Scherer, and G. Shan, “Evaluation of logistic and polynomial models for fitting sandwich-ELISA calibration curves,” J Immunol Methods, vol. 339, no. 2, pp. 245–258, Dec. 2008, doi: 10.1016/J.JIM.2008.09.001.

[45] L. Zhang et al., “Proteomic Analysis of Vesicle-Producing Pseudomonas aeruginosa PAO1 Exposed to X-Ray Irradiation,” Front Microbiol, vol. 11, Dec. 2020, doi: 10.3389/fmicb.2020.558233.

[46] Z. Ayed, L. Cuvillier, G. Dobhal, and R. V. Goreham, “Electroporation of outer membrane vesicles derived from Pseudomonas aeruginosa with gold nanoparticles,” SN Appl Sci, vol. 1, no. 12, p. 1600, Dec. 2019, doi: 10.1007/s42452-019-1646-2.

[47] M. Paulsson et al., “Peptidoglycan-Binding Anchor Is a Pseudomonas aeruginosa OmpA Family Lipoprotein With Importance for Outer Membrane Vesicles, Biofilms, and the Periplasmic Shape,” Front Microbiol, vol. 12, Feb. 2021, doi: 10.3389/fmicb.2021.639582.

[48] S.-W. Hong et al., “Extracellular vesicles derived from Staphylococcus aureus induce atopic dermatitis-like skin inflammation,” Allergy, vol. 66, no. 3, pp. 351–359, Mar. 2011, doi: 10.1111/j.1398-9995.2010.02483.x.

[49] B. S. R. da Luz et al., “Impact of Environmental Conditions on the Protein Content of Staphylococcus aureus and Its Derived Extracellular Vesicles,” Microorganisms, vol. 10, no. 9, p. 1808, Sep. 2022, doi: 10.3390/microorganisms10091808.

[50] P. Deo et al., “Mitochondrial dysfunction caused by outer membrane vesicles from Gram-negative bacteria activates intrinsic apoptosis and inflammation,” Nat Microbiol, vol. 5, no. 11, pp. 1418–1427, Nov. 2020, doi: 10.1038/s41564-020-0773-2.

[51] N. J. Bitto et al., “Bacterial membrane vesicles transport their DNA cargo into host cells,” Sci Rep, vol. 7, no. 1, Dec. 2017, doi: 10.1038/s41598-017-07288-4.

[52] M. Hadadi-Fishani, S. Najar-Peerayeh, S. Davar Siadat, M. Sekhavati, and A. Mohabati Mobarez, “Isolation and immunogenicity of extracted outer membrane vesicles from Pseudomonas aeruginosa under antibiotics treatment conditions,” 2021. [Online]. Available: http://ijm.tums.ac.ir

[53] N. Couto, S. R. Schooling, J. R. Dutcher, and J. Barber, “Proteome Profiles of Outer Membrane Vesicles and Extracellular Matrix of Pseudomonas aeruginosa Biofilms,” J Proteome Res, vol. 14, no. 10, pp. 4207–4222, Oct. 2015, doi: 10.1021/acs.jproteome.5b00312.

[54] C. Pérez-Cruz, L. Delgado, C. López-Iglesias, and E. Mercade, “Outer-Inner Membrane Vesicles Naturally Secreted by Gram-Negative Pathogenic Bacteria,” PLoS One, vol. 10, no. 1, p. e0116896, Jan. 2015, doi: 10.1371/journal.pone.0116896.

[55] M. M. E. Metruccio, D. J. Evans, M. M. Gabriel, J. L. Kadurugamuwa, and S. M. J. Fleiszig, “Pseudomonas aeruginosa Outer Membrane Vesicles Triggered by Human Mucosal Fluid and Lysozyme Can Prime Host Tissue Surfaces for Bacterial Adhesion,” Front Microbiol, vol. 7, Jun. 2016, doi: 10.3389/fmicb.2016.00871.

[56] F. Y. McWhorter, T. Wang, P. Nguyen, T. Chung, and W. F. Liu, “Modulation of macrophage phenotype by cell shape,” Proceedings of the National Academy of Sciences, vol. 110, no. 43, pp. 17253–17258, Oct. 2013, doi: 10.1073/pnas.1308887110.

[57] J. C. Santos et al., “LPS targets host guanylate-binding proteins to the bacterial outer membrane for non-canonical inflammasome activation,” EMBO J, vol. 37, no. 6, p. e98089, Mar. 2018, doi: 10.15252/embj.201798089.

[58] A. J. Park et al., “Tracking the dynamic relationship between cellular systems and extracellular subproteomes in Pseudomonas aeruginosa biofilms,” J Proteome Res, vol. 14, no. 11, pp. 4524–4537, Nov. 2015, doi: 10.1021/acs.jproteome.5b00262.

[59] I. M. Torres, S. Demirdjian, J. Vargas, B. C. Goodale, and B. Berwin, “Acidosis increases the susceptibility of respiratory epithelial cells to Pseudomonas aeruginosa -induced cytotoxicity,” American Journal of Physiology-Lung Cellular and Molecular Physiology, vol. 313, no. 1, pp. L126–L137, Jul. 2017, doi: 10.1152/ajplung.00524.2016.

[60] F. Dell’Annunziata et al., “Klebsiella pneumoniae-OMVs activate death-signaling pathways in Human Bronchial Epithelial Host Cells (BEAS-2B),” Heliyon, vol. 10, no. 8, p. e29017, Apr. 2024, doi: 10.1016/j.heliyon.2024.e29017.

[61] S. Santos-Pereira, F. O. Cardoso, K. S. Calabrese, and T. Zaverucha do Valle, “Leishmania amazonensis resistance in murine macrophages: Analysis of possible mechanisms,” PLoS One, vol. 14, no. 12, p. e0226837, Dec. 2019, doi: 10.1371/journal.pone.0226837.

[62] S. B. Corradin and J. Mauël, “Phagocytosis of Leishmania enhances macrophage activation by IFN-gamma and lipopolysaccharide.,” J Immunol, vol. 146, no. 1, pp. 279–285, Jan. 1991.

[63] C. Schwechheimer and M. J. Kuehn, “Outer-membrane vesicles from Gram-negative bacteria: biogenesis and functions,” Nat Rev Microbiol, vol. 13, no. 10, pp. 605–619, Oct. 2015, doi: 10.1038/nrmicro3525.

[64] H. Bichiou, C. Bouabid, I. Rabhi, and L. Guizani-Tabbane, “Transcription Factors Interplay Orchestrates the Immune-Metabolic Response of Leishmania Infected Macrophages,” Front Cell Infect Microbiol, vol. 11, Apr. 2021, doi: 10.3389/fcimb.2021.660415.

[65] Z. Xu, X. Hao, M. Li, and H. Luo, “Rhodococcus equi-Derived Extracellular Vesicles Promoting Inflammatory Response in Macrophage through TLR2-NF-κB/MAPK Pathways,” Int J Mol Sci, vol. 23, no. 17, p. 9742, Aug. 2022, doi: 10.3390/ijms23179742.

[66] T. F. Murphy et al., “Pseudomonas aeruginosa in Chronic Obstructive Pulmonary Disease,” Am J Respir Crit Care Med, vol. 177, no. 8, pp. 853–860, Apr. 2008, doi: 10.1164/rccm.200709-1413OC.

[67] J. I. Weber et al., “Insights on Host–Parasite Immunomodulation Mediated by Extracellular Vesicles of Cutaneous Leishmania shawi and Leishmania guyanensis,” Cells, vol. 12, no. 8, p. 1101, Apr. 2023, doi: 10.3390/cells12081101.

[68] L. E. Emerson et al., “Leishmania infection-derived extracellular vesicles drive transcription of genes involved in M2 polarization,” Front Cell Infect Microbiol, vol. 12, Aug. 2022, doi: 10.3389/fcimb.2022.934611.

