## Supplementary figures and images for "Impact of Bacterial Membrane Vesicles on Cellular Responses in *Leishmania amazonensis*-Infected Macrophages *In Vitro*"

### Supplementary Figure 1

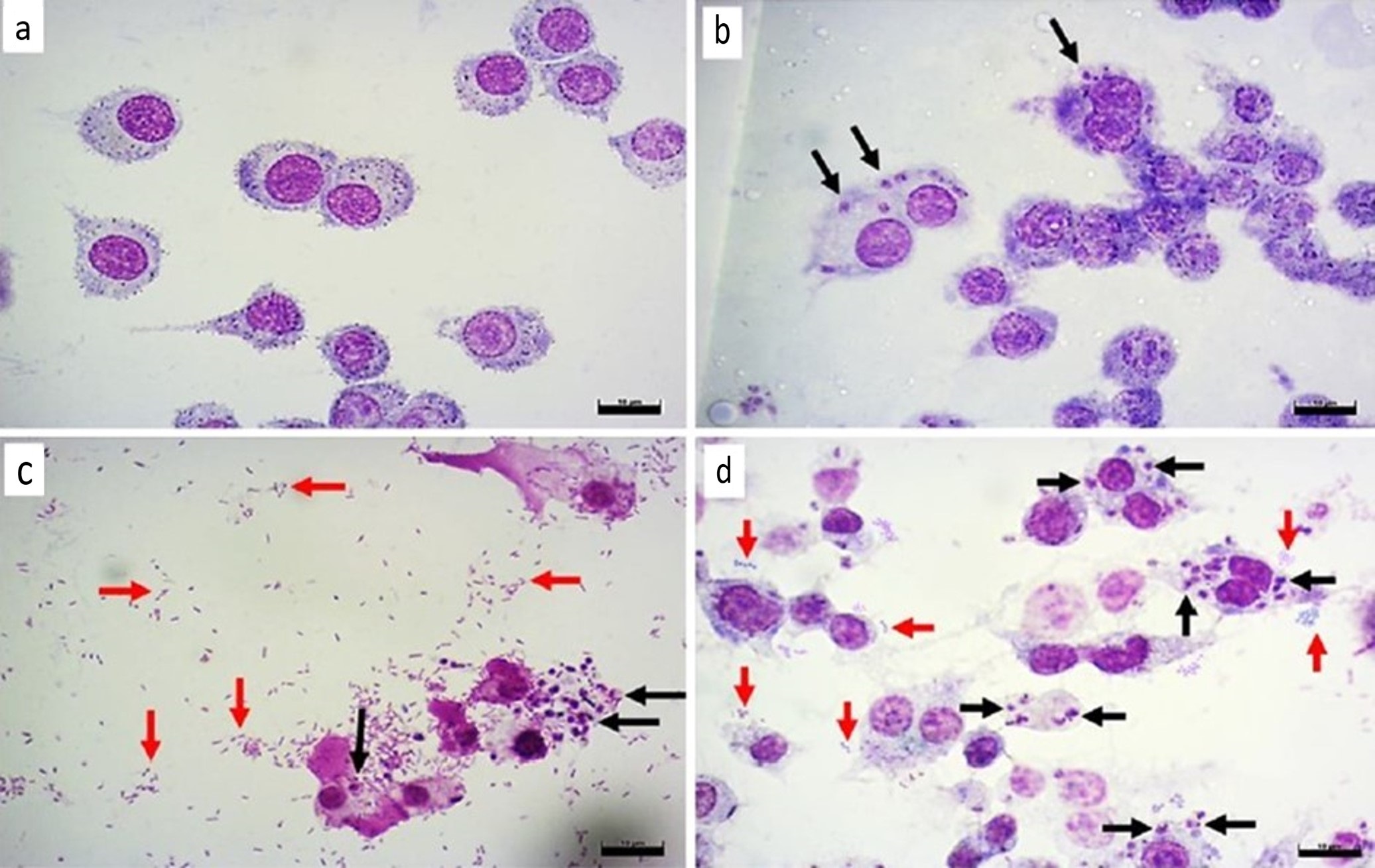
